# Molecular determinants underlying differential recruitment of p115RhoGEF and PDZRhoGEF to activated Gα_13_

**DOI:** 10.1101/2024.12.28.630591

**Authors:** Anna Fassler Bakhman, Viktoriya Lukasheva, Christian Le Gouill, Mickey Kosloff, Michel Bouvier

**Author notes:** Correspondence to: Dr. Mickey Kosloff, Department of Human Biology, Faculty of Natural Sciences, University of Haifa, 199 Aba Khoushy Ave., POB 3338, Haifa, 3103301, Israel. And Dr. Michel Bouvier, Department of Biochemistry and Molecular Medicine & Institute for Research in Immunology and Cancer, Université de Montréal, 2900, Boulevard Édouard-Montpetit, Pavillon Marcelle Coutu, Montréal, Qc., H3T 1J4, Canada.

## Abstract

Heterotrimeric G proteins, particularly Gα_12_ and Gα_13_, are pivotal regulators of cellular signaling pathways. Their direct downstream effectors, which include p115RhoGEF and PDZRhoGEF, engage downstream signaling via Rho activation. Yet the molecular determinants that dictate their differential recruitment by Gα_12/13_ are not fully understood. Here, we combined quantitative computational residue-level analysis with site-directed mutagenesis and bioluminescence resonance energy transfer (BRET)-based assays to dissect Gα_13_ interactions with these RhoGEFs. We mapped the contributions of individual residues to binding and identified specific Gα_13_ residues in its helical domain, switch regions, and effector-binding site as key yet differential contributors to p115RhoGEF and PDZRhoGEF recruitment. Experimental validation with BRET confirmed that changes in many Gα_13_ residues impact p115RhoGEF more substantially than PDZRhoGEF, underscoring the specificity of Gα_13_ interactions with p115RhoGEF. Investigation of the p115RhoGEFs identified critical residues that contribute to interactions with Gα_13_ and Gα_12_. Our findings highlight residue-level differences in the molecular interactions of Gα_13_ with p115RhoGEF and PDZRhoGEF, providing insights into the specificity and regulation of Gα_13_-mediated signaling pathways. The resulting residue-level maps lay the groundwork for development of selective therapeutic strategies targeting Gα_13_-RhoGEF interactions.

## INTRODUCTION

Heterotrimeric (αβγ) G proteins are key regulators of cellular signaling pathways, orchestrating a wide variety of responses within the cell. Among the 16 human genes that encode for Gα subunits, GNA12 and GNA13 encode for the Gα_12_ and Gα_13_ subunits, which are < 45% identical in sequence to other Gα subunits but share 67% amino acid identity with each other (1). Gα_12_ and Gα_13_ (Gα_12/13_) are key regulators of cytoskeletal dynamics and mediate a variety of physiological processes, including embryonic development, cell growth and migration, angiogenesis, platelet activation, immune response, apoptosis, and neuronal responses (2-10). Abnormal regulation of Gα_12/13_-related pathways has been found in pathologies such as oncogenesis, tumor cell invasion and metastasis as well as hypertension (11-16). Gα_12/13_, like all heterotrimeric G proteins, are activated by G protein-coupled receptors (GPCRs), leading to exchange of GDP for GTP in the Gα active site, while Gα inactivation is regulated by GTPase activating proteins (GAPs) that accelerate GTP hydrolysis (17-20). The activated Gα_12/13_ proceeds to activate downstream signaling mainly via proteins from the RhoGEF family (19, 21-25). Specifically, three RhoGEFs – p115RhoGEF, PDZRhoGEF, and leukemia-associated RhoGEF (LARG) – are directly modulated by the Gα_12/13_ subfamily, with p115RhoGEF also having reciprocal GAP activity, enhancing Gα_12/13_ inactivation (22, 23, 26-30). However, the specific residue-level determinants that govern Gα_12/13_-p115/PDZRhoGEF interactions have yet to be fully elucidated.

Structural and biochemical studies mostly used Gα_13_ as a representative of the Gα_12/13_ family and showed that Gα_13_ interacts with p115RhoGEF and PDZRhoGEF via both the Gα “helical domain” and the Gα “GTPase domain” (31-33). The structure of the Gα_13_-p115RhoGEF complex demonstrated that the Gα_13_ GTPase domain was at the heart of this complex and provided most of its interaction interface with p115RhoGEF (31). The Gα_13_ residues participating in this interface were located mostly in the three Gα_13_ switch regions (Sw I-III), which change their conformation depending on whether the Gα subunit binds GTP or GDP (31, 32, 34), and in the Gα_13_ “effector binding region”, which encompasses the C-terminus of Sw III, the α3 helix, and the following α3-β5 loop (31-33, 35, 36). The importance of these Gα regions in interactions with RhoGEFs was emphasized by mutagenesis studies. Charge reversal mutations in the Gα_13_ Sw I (K204E) and Sw II (E229K or R232E) completely abolished interactions with p115RhoGEF (37). Mutagenesis of Gα_12_ also showed the importance of this domain, as deletion mutations of six residues in the C-terminus of the Gα_12_ Sw II resulted in a complete loss of p115RhoGEF binding (38). Mutations within the Gα_13_ effector binding region (T274E, N278A, and T274E/N278A) also impaired binding to p115RhoGEF and affected its GAP activity – reducing it by 50% (N278A) or entirely (T274E and T274/N278) (33). Nevertheless, the interactions of Gα_13_ with p115RhoGEF and PDZRhoGEF have mostly been investigated in separate studies, while direct comparisons reveal specific differences in how Gα_13_ binds these RhoGEFs are lacking.

On the RhoGEF side of the interface, interactions with Gα_12/13_ have been shown to occur via both the RhoGEF N-terminal domain (NTD) and its RGS homology (RH) domain (31-33). The NTD, which is approximately 20 amino acids long, precedes the 180 amino acids long RH domain. All three RhoGEFs also contain Dbl homology (DH) and Pleckstrin homology (PH) domains that mediate Rho activation and GEF activity. While the DH domain was shown to interacts with Gα_13,_ in a position opposite to the binding site of the NTD and RH domains, the PH domain was not seen to interact with Gα_13_ (39). PDZRhoGEF also contains an additional N-terminal PDZ domain that, similar to the PH domain, does not directly interact with Gα_13_ (40) and is absent from p115ThoGRF and LARG. The NTD and the RH domain engage with both the GTPase domain and the helical domain of Gα_13_ (31-33). Mutagenesis of the p115RhoGEF NTD showed that it is necessary for p115RhoGEF GAP activity (31, 32). Specifically, p115RhoGEF residues E27-E29 and F31 were shown as crucial both for binding Gα_13_ and for p115RhoGEF GAP activity (36). Similarly, mutations in the ^308^IIG motif in the PDZRhoGEF NTD, which is conserved across all RH-containing RhoGEFs and corresponds to the ^23^IIG motif in p115RhoGEF, showed this motif was vital for interactions with Gα_12/13_, particularly with Sw I (32). However, the absence of GAP activity for PDZRhoGEF was attributed to differences in the NTD between the RhoGEFs (30, 33, 36, 41). Overall, these previous studies highlighted the importance of both the p115RhoGEF NTD and RH domain for GAP activity, yet the root for differences in interactions between p115RhoGEF and PDZRhoGEF with Gα_13_ are less clear.

In this study, we employed energy-based calculations with experimentally determined structures to produce a quantitative residue-level map of Gα_13_ interactions with p115RhoGEF and PDZRhoGEF, enabling a detailed comparison of Gα_13_ interactions with the two RhoGEFs. We combined these computational results with site-directed mutagenesis and bioluminescence resonance energy transfer (BRET) sensors using G protein effector membrane translocation assays (GEMTA) (42), based on enhanced bystander BRET (ebBRET) (43), which enable to precisely measure the effect of mutations on p115RhoGEF and PDZRhoGEF recruitment. Thereby, we interrogated the residue-level determinants of Gα_12/13_-RhoGEF binding and identified key residues on both sides of the interface, demonstrating essential differences in Gα_13_ interactions with p115RhoGEF and PDZRhoGEF.

## RESULTS

### Residue-level computational mapping of Gα*_13_* interactions with RhoGEFs

To map the individual residues that contribute to the interactions of Gα_13_ with p115RhoGEF and PDZRhoGEF, we used energy calculations to analyze the following four structures of Gα_13_ with p115RhoGEF and with PDZRhoGEF: two structures of GDP/AlF_4_-bound Gα_13_ with p115RhoGEF (PDB ID 3AB3 and 1SHZ), GDP/AlF_4_-bound Gα_13_ with PDZRhoGEF (PDB ID 3CX7), and GTPγS-bound Gα_13_ with PDZRhoGEF (PDB ID 3CX8) (31-33). Following previous work, we calculated polar/electrostatic contributions separately from non-polar/hydrophobic interactions. Across the four complexes detailed above, we used the Finite Difference Poisson–Boltzmann (FDPB) method to calculate the net electrostatic and polar contributions (ΔΔG_elec_) of each residue within 15 Å of the Gα_13_-RhoGEF interface. For each residue, we separately calculated the electrostatic contributions from the side chain and/or those originating from the main chain of each residue. Residues that substantially contribute electrostatically to the interaction were defined as those contributing ΔΔG_elec_ ≥ 1 kcal/mol, i.e., twice the maximal numerical error of the electrostatic calculations (44). This approach calculates the net difference between the electrostatic interaction of a residue with its protein partner in relation to its interaction with the water and ions in the solvent, and thereby identifies only residues that are calculated to substantially contribute to binding. In parallel, non-polar energy contributions (ΔΔG_np_) for each residue were calculated as a surface-area proportional term by multiplying the per-residue surface area buried upon complex formation by a surface tension constant of 0.05 kcal/mol/Å^2^. Residues with substantial non-polar contributions were defined as those contributing ΔΔG_np_ ≥ 0.5 kcal/mol to the interactions (namely, more than 10Å^2^ of each protein surface is buried upon complex formation). To reduce false negatives that come from non-representative side chain rotamers in an individual structural snapshot, we applied a consensus approach by comparing biological “replicates” across comparable PDB structures (Figs. S1 and S2). When analyzing the two Gα_13_-PDZRhoGEF complexes with GTPγS and GDP/AlF_4_, we combined the results from either complex into a consensus approach that unified both sets of predictions (see Materials and Methods for details). We compared the calculated energies both in the presence and absence of the nucleotide, observing no significant effect on the results, as expected due to the fact that the nucleotide buried within the Gα subunit is distant from the interface with RhoGEFs. Residues calculated to contribute substantially to intermolecular interactions were mapped onto the 3D structure of each individual protein (Fig. 1).

**Figure 1.**
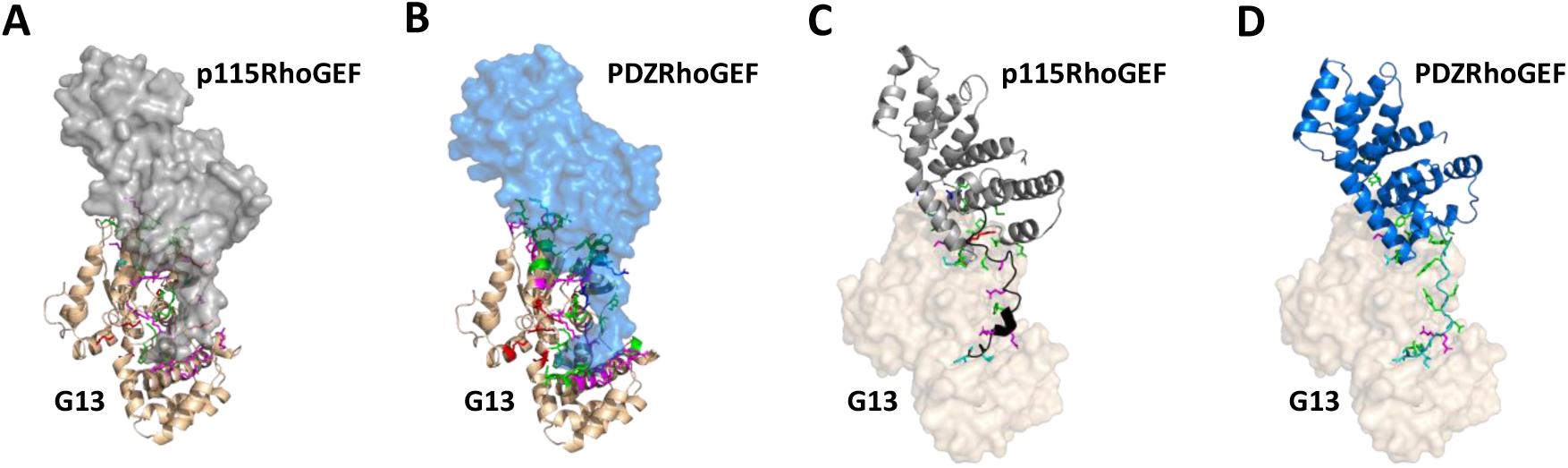
Residues that contribute substantially to interactions in Gα_13_ complexes with p115RhoGEF and PDZRhoGEF. **A**. Residues in Gα_13_ (wheat ribbon) that contribute to interactions with p115RhoGEF (gray molecular surface), shown as sticks and colored green (non-polar contributions), red (side-chain electrostatic contributions), cyan (non-polar and main-chain electrostatic contributions), blue (main-chain electrostatic contributions), and magenta (non-polar and side-chain electrostatic contributions). **B**. Residues in Gα_13_ that contribute to interactions with PDZRhoGEF (blue molecular surface), colored as in A. **C**. Residues in p115RhoGEF that contribute to interactions with Gα_13_ (wheat molecular surface), shown as sticks and colored as in A. p115RhoGEF is shown as a ribbon diagram colored black (NTD) and gray (RH domain). **D**. Residues in PDZRhoGEF that contribute to interactions with Gα_13_, shown as sticks and colored as in B. PDZRhoGEF is shown as a ribbon diagram colored teal (NTD) and blue (RH domain).

Our analysis identified 43 and 45 Gα_13_ residues that contribute substantially to interactions with p115RhoGEF and PDZRhoGEF, respectively (Figs. 1A-B, S1, S2). Of these contributing residues, ∼25-30% are located at the Gα_13_ helical domain and the rest in the GTPase domain. Out of the contributing GTPase domain residues, more than half are in the three switch regions and ∼40% are located within the effector binding region. The remaining small minority of contributing residues are located in the Gα_13_ N-terminal α1 helix and in the C-terminal region.

On the opposing side of the interface, our calculations identified 32 p115RhoGEF residues and 34 PDZRhoGEF residues that contribute to the interactions with Gα_13_ (Figs. 1C-D, S4, S5). The two RhoGEFs share only 33% sequence identity and have no significant sequence similarity between their NTDs. Nevertheless, in both p115RhoGEF and PDZRhoGEF ∼40% of the contributing residues are located in the NTD. The contributions that come from the RH domains of both RhoGEFs originate from three distinct regions that, nevertheless, combine together when interacting with Gα_13_ (Fig. 1C-D). Interestingly, the number of RhoGEF residues that contribute electrostatically is ∼2/3 of the corresponding residues that contribute electrostatically from the Gα_13_ side (Figs. 2, S4, S5). Visual inspection showed that in several cases, this apparent discrepancy comes from two or three Gα_13_ residues interacting electrostatically with a single RhoGEF residue. These asymmetric electrostatic interactions with Gα_13_ are found across both RhoGEFs and involve mainly NTD residues.

**Figure 2.**
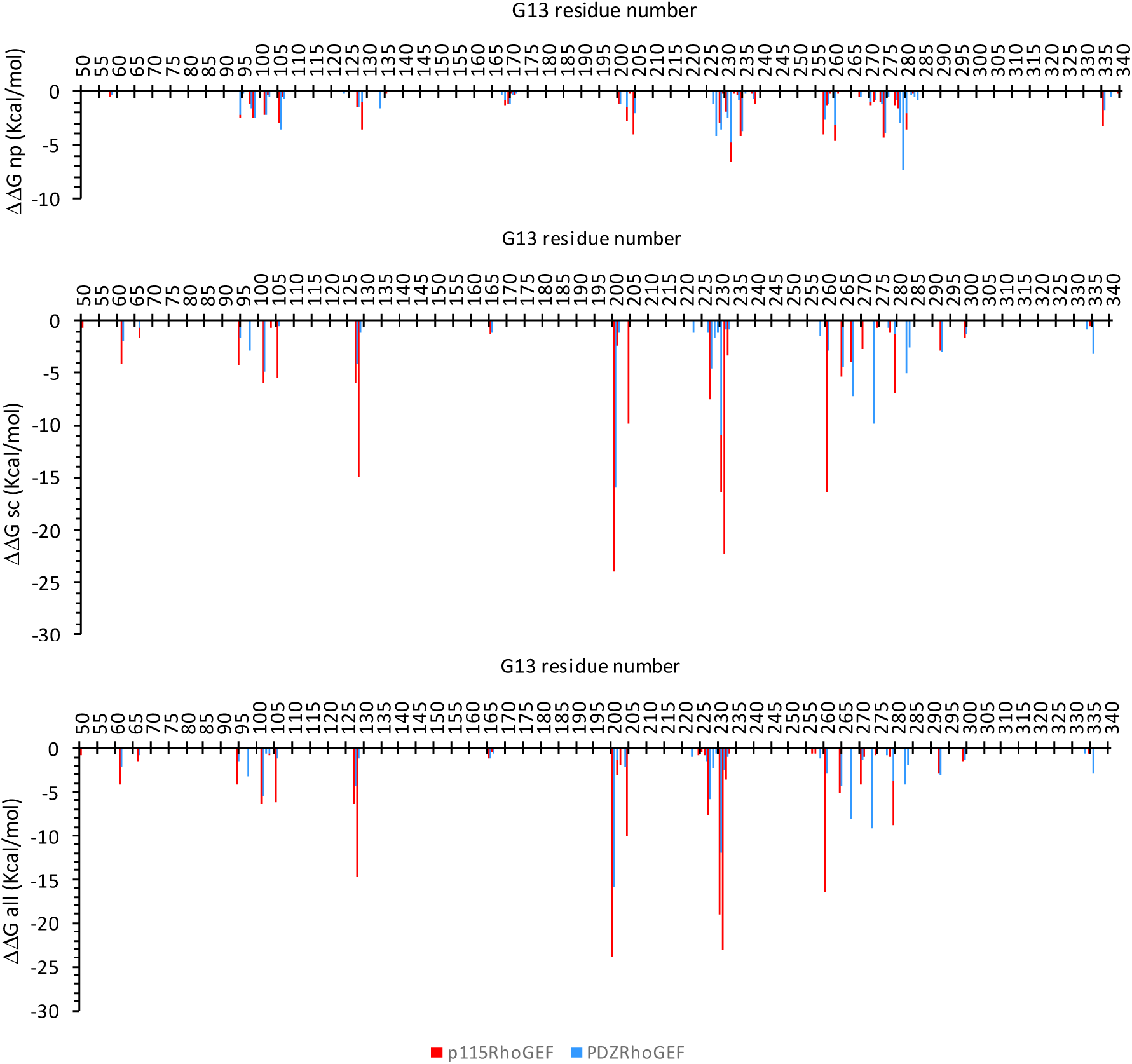
Gα_13_ per-residue energy contributions to interactions with p115RhoGEF and with PDZRhoGEF. Panels show the energy results for non-polar (np) contributions, electrostatic contributions from the side-chain (sc), and electrostatic contributions from the entire residue (all), calculated as described in Methods for the Gα_13_-p115RhoGEF complexes (PDB ID 3AB3 and 1SHZ) and the Gα_13_-PDZRhoGEF complexes (PDB ID 3CX7 and 3CX8) and combined into one graph using a consensus approach.

**Figure 3.**
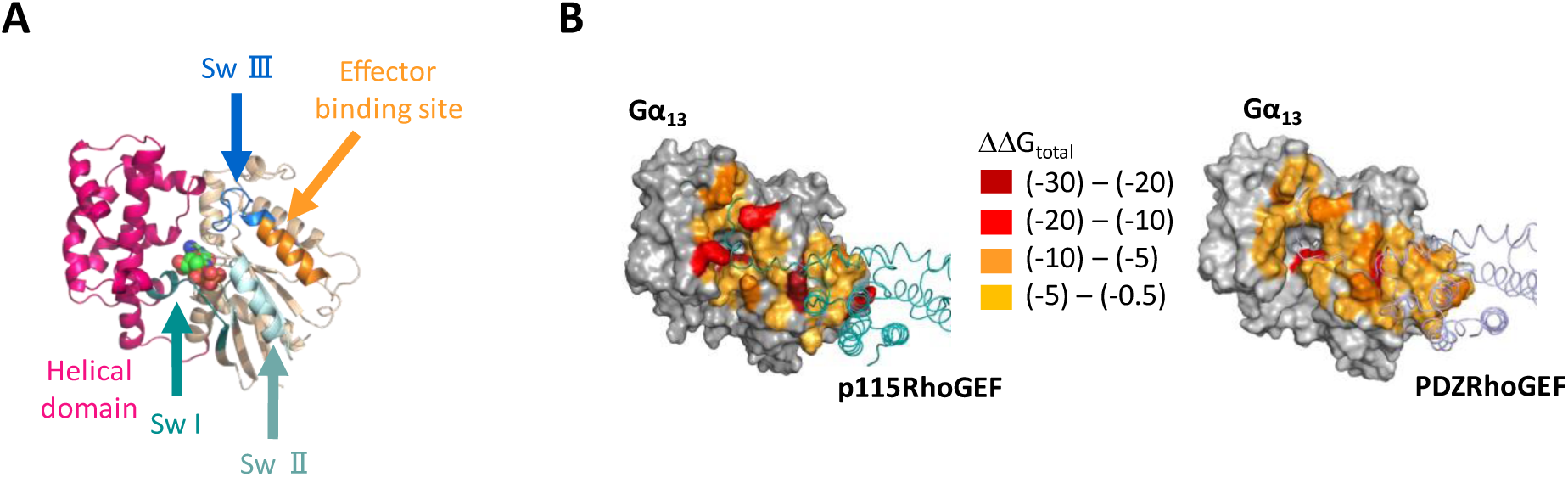
Residues in Gα_13_ contribute more substantially to interactions with p115RhoGEF than to interactions with PDZRhoGEF. **A.** Structural regions in Gα_13_ that can interact with its partners. Gα_13_ is shown as a ribbon diagram colored wheat (GTPase domain) and magenta (helical domain). The nucleotide is shown as spheres, colored by element. **B.** Total per residue energy contributions (ΔΔGtotal), which equal the sum of the non-polar and electrostatic contributions of Gα_13_ (Fig. 2) to p115RhoGEF (left) and PDZRhoGEF (right), as calculated in Fig. S3.

**Figure 4.**
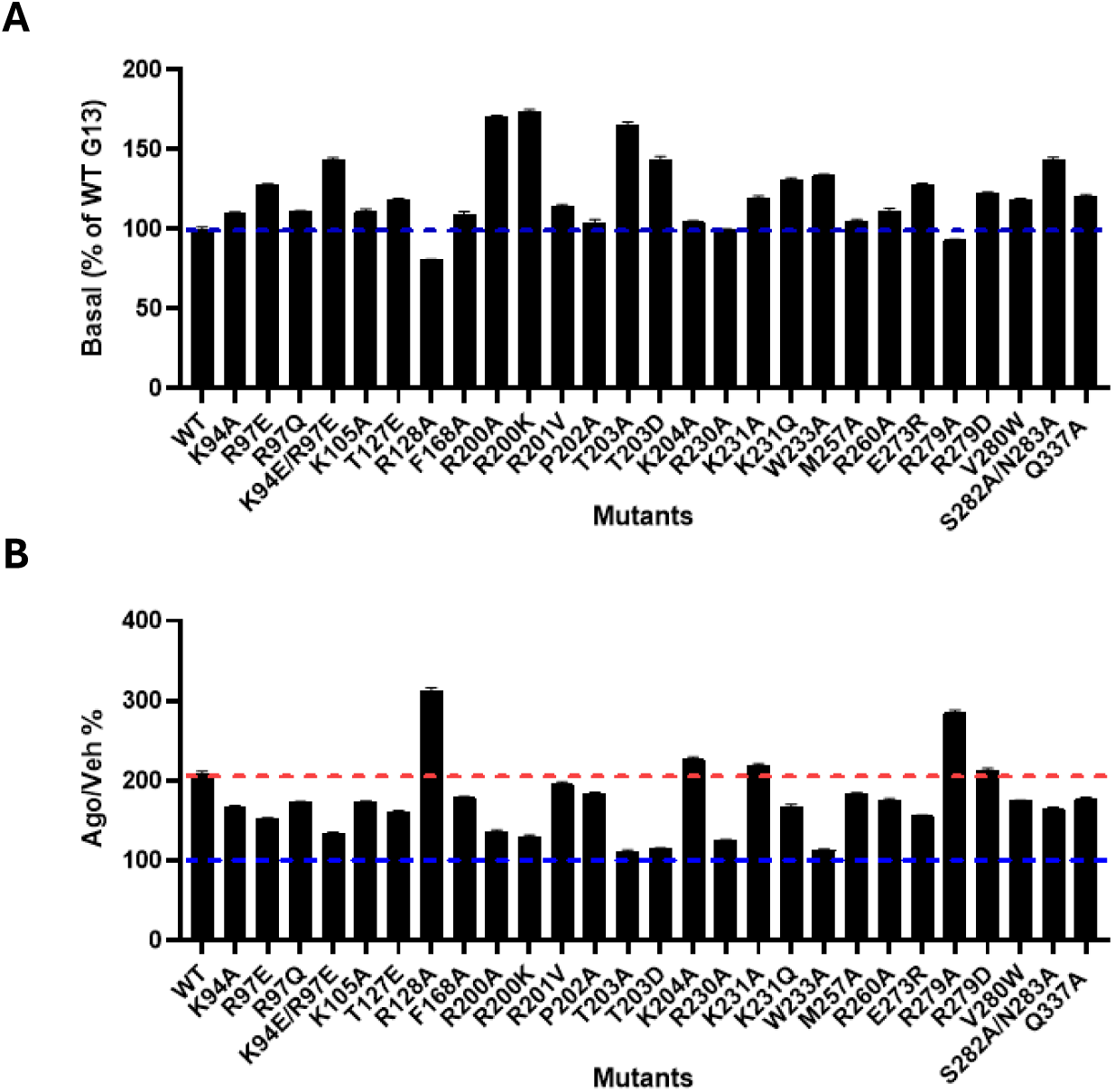
Effect of Gα_13_ mutations on basal (A) and TPαR-promoted activation (B) of Gα_13_ assessed using a GRK-Gβγ-based sensor monitoring the release of free Gβγ from WT and mutant Gα_13_. BRET was measured between GRK2-D110-GFP10 and Gγ5-Rluc in the presence of heterologously expressed wild-type (WT) or mutant forms of Gα_13_ under basal condition (A) or following activation of the thromboxane receptor TPα with U46619 (B). **A.** The basal signal for each mutant is expressed as % of the signal observed for WT Gα_13_. **B.** The U46619-promoted response is expressed as % of the vehicle. 100% (the solid line) indicates the value observed in the absence of stimulation (vehicle) for WT-Gα_13_. The dotted line indicates U46619-promoted response for the WT-Gα_13_.

**Figure 5.**
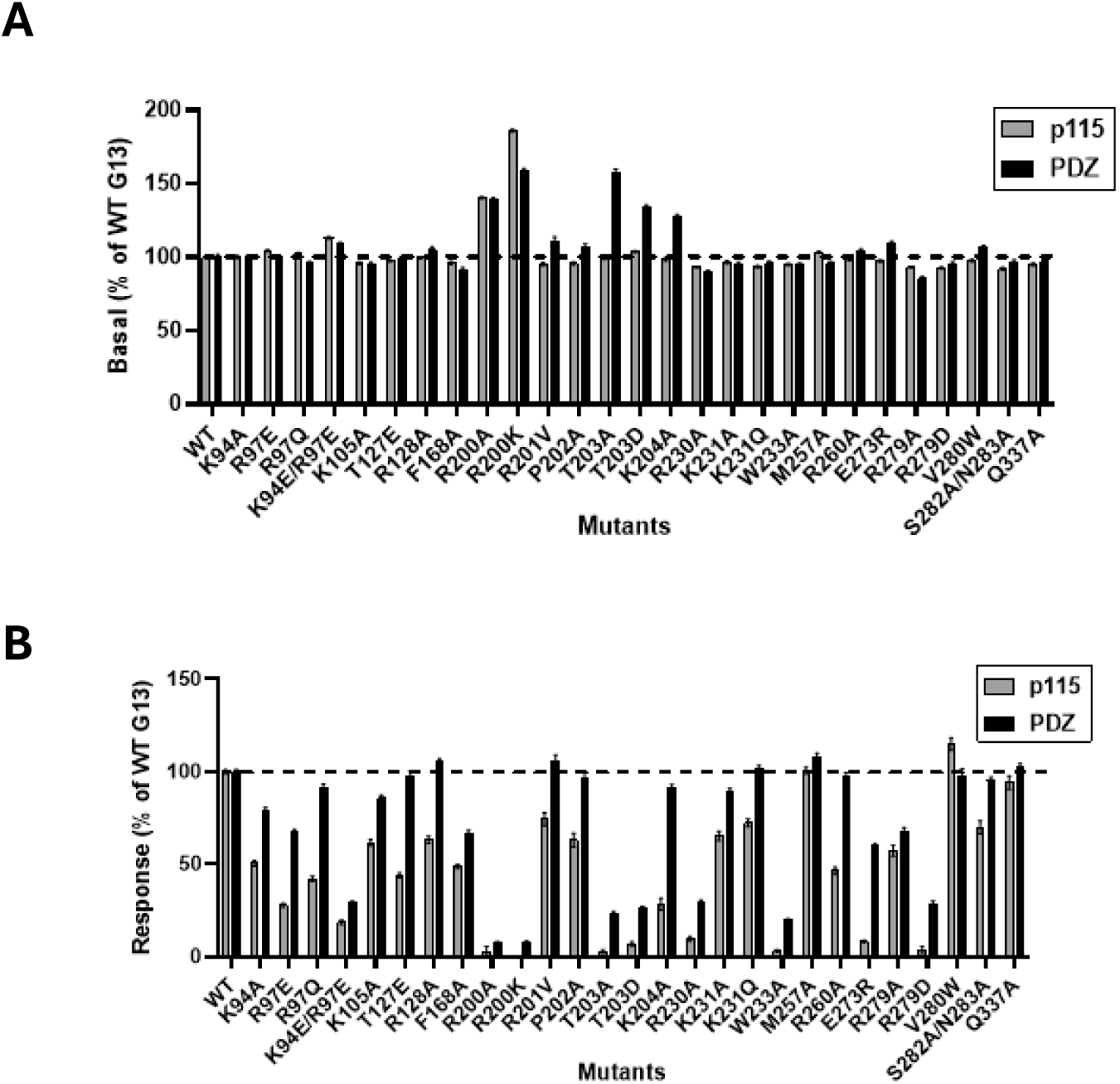
Effect of Gα_13_ mutations on basal (A) and TPαR-promoted recruitment (B) of p115RhoGEF and PDZRhoGEF to Gα_13_ assessed using GEMTA sensors. BRET was measured between the plasma-anchored rGFP-CAAX and either p115-RhoGEF-Rluc (p115; blue bars) or PDZRhoGEF-Rlucz (PDZ; red bars) in the presence of heterologously expressed WT or mutant forms of Gα_13_ under basal condition (A) or following activation of the thromboxane receptor TPα with U46619 (B). **A.** The basal signal for each mutant is expressed as % of the signal observed for wild-type (WT) Gα_13_. **B.** The U46619-promoted response is expressed as % of the response observed for WT Gα_13_ shown as 100% (the dotted line).

The Gα_13_ positions that contribute to interactions with both RhoGEFs are mostly identical – 39 Gα_13_ residues contribute to both p115RhoGEF and PDZRhoGEF (Fig. 2). The few Gα residues that contribute to only one RhoGEF do so via weaker and usually non-polar contributions. However, in the vast majority of the Gα_13_ positions, across all Gα_13_ regions, contributions to p115RhoGEF were substantially stronger than to PDZRhoGEF (Fig. 2). This effect was especially pronounced in electrostatic contributions, with a majority of Gα_13_ interactions to p115RhoGEF showing ∼1.5-3 fold higher energy values than those for PDZRhoGEF. Comparing the total energy contributions (ΔΔG_total_, which equal to the sum of ΔΔG_elec_ and ΔΔG_np_) we observed that these quantitative differences were especially prominent in Gα_13_ helical domain and Sw II residues (Figs. 3, S3). Two particular Gα_13_ residues, R128 (helical domain) and K231 (Sw II), which interact with the NTD of both RhoGEFs, stood out; the energy contributions of these two Gα_13_ residues to p115RhoGEF were 4-fold and 17-fold stronger than to PDZRhoGEF, respectively.

### Mutations across Gα_13_ disrupt recruitment of both RhoGEFs but with a stronger effect on p115RhoGEF

Our computational residue-level analysis pinpointed seven Gα_13_ positions as key contributors to interactions with p115RhoGEF and PDZRhoGEF, with a ΔΔG_total_ of more than -10 Kcal/mol each (Figs. 3, S3). To test the importance of these Gα_13_ residues for RhoGEF binding, we used site-directed mutagenesis and ebBRET assays to quantify Gα_13_–RhoGEF interactions with G protein effector membrane translocation assay (GEMTA) biosensors (42, 45). These biosensors consist of the NTD and RH domain of p115RhoGEF and PDZRhoGEF, (p115RhoGEF 21-244 and PDZRhoGEF 281-483) fused to *Renilla* luciferase (RlucII) following their respective RH domain. Activation of the GPCR leads to Gα_12/13_-dependent recruitment of the RlucII-fused effectors to the plasma membrane, bringing the RlucII energy donor near the energy acceptor, *Renilla* green fluorescent protein, that is anchored to the plasma membrane through a CAAX motif derived from K-Ras (rGFP-CAAX, Fig. S7). Thus, RhoGEF binding to activated Gα_12/13_ leads to an increase in the BRET signal (42, 45). The advantage of such assay design is that it does not require the modification of the G protein subunits. The impact of mutations on engagement with RhoGEFs was assessed upon Gα_13_ activation by the thromboxane receptor TPα, a receptor known to strongly activate Gα_13_ (46).

In order to assess whether the mutations could affect the activation of the G protein *per se,* we monitored the impact of the mutations on the dissociation of Gα and Gβγ using a GRK-based, sensor monitoring the release of free Gβγ from WT and mutant forms of Gα (47) (Fig. S7). As can be seen in Fig. 4A, several mutations (K94E/R97E, R200A, R200K, T203A, T203D and S282A/N283), which are in residues that are not in the Gα-Gβγ interface, led to constitutive free Gβγ. This may indicate either a reduced affinity of the mutant Gα for Gβγ or a spontaneous release of Gβγ that would reflect a level of constitutive activity of the Gα mutant. We note that R200 is the catalytic arginine that interacts directly with the nucleotide and, therefore, mutation of this residue directly affects activation and dissociation from Gβγ. Consistent with the previous observation that PKA phosphorylation of Gα-T203 destabilizes coupling to Gβγ (48), it is interesting to note that mutation of T203 to either A or D led to a strong constitutive free Gβγ. This residue, while not at the Gα interface with Gβγ, is located between K204 and H207, both of which do interact with Gβγ. This positioning suggests that upon mutation T203 indirectly affects interactions with Gβγ by changing the conformation of these two Gα residues.

Upon activation with TPα using U46619 as the agonist, only two mutants showed increased activation (R128A, R279A), whereas many mutants showed partial reduction in activation and only three (T203A, T203D, and W233A) showed no activation at all (Fig. 4B). The lack of detectable activation for these three mutants most likely results from elevated basal signal, resulting from constitutive activity or reduced affinity for Gβγ. Notably, W233 is directly at the interface with Gβγ, which explains the decreased affinity to Gβγ upon mutation of this residue. Altogether, these data show that most mutants can be activated by a GPCR and thus are amenable for comparing their ability to recruit p115RhoGEF vs PDZ-RhoGEF.

When considering the impact of the mutations on the recruitment of the p115RhoGEF and PDZRhoGEF biosensors, we first assessed the effects on the constitutive recruitment of the RhoGEFs by the mutants. Notable basal recruitment of five mutants was observed – R200A, R200K, T203A, T203D, K204A (Fig. 5A). Interestingly, mutations of T203A, T203D, and K204A only promoted constitutive recruitment for PDZRhoGEF and not p115RhoGEF. Given the constitutive recruitment of PDZRhoGEF by the T203 mutants, which is consistent with the increased basal free Gβγ observed in Fig. 4A for T203A and T203D, the lack of such constitutive recruitment of p115RhoGEF could results from a direct impact of the mutation on the Gα_13_-p115RhoGEF interaction. Additionally, K204, which is at the interface with both Gβγ and RhoGEFs, contributes more substantially to interactions with p115RhoGEF than with PDZRhoGEF (Fig. 2). This suggests that the K204A

Upon stimulation with TPα, among the mutations selected for their predicted effect on RhoGEF interaction, most affected p115RhoGEF more than PDZRhoGEF (Fig. 5B). The impact of the mutations on the maximal p115RhoGEF and PDZRhoGEF recruitment promoted by the TPα agonist and its potency is presented in Table S1. Only three mutations did not affect either RhoGEFs (M257A, V280W. and Q337A). Interestingly, some mutations selectively affected p115RhoGEF over PDZRhoGEF. These include the R97Q, K105A, T127E, R128A, R201C, P202A, K204A, K231A,

K231Q, S282A/N283A mutants that were found to affect only p115RhoGEF. This aligns with our computational analysis, as most residues contribute more substantially to Gα_13_-p115RhoGEF interactions than to interactions with PDZRhoGEF, while the remainder of these residues contribute only to interactions with p115RhoGEF. The only exception is the S282/N293 mutant, as S282 and N293 were predicted to contribute only to interactions with PDZRhoGEF. In addition, some mutants that could not recruit p115RhoGEF, maintain a measurable ability to recruit PDZRhoGEF (T203A/D, R230A, W233A, E273R and R279D), which is also in line with our computational results. While T203 does not contribute directly to interactions with p115RhoGEF, it is located between R201 and K204, which contribute substantially only to interactions with p115RhoGEF, and can affect these interactions upon mutation. Additionally, R230, E273, and R279 make a substantially larger contribution to interactions with p115RhoGEF than with PDZRhoGEF. In contrast with the R279D mutation, substitution to an alanine at this position (R279A) results in a 50% reduction in recruitment of both RhoGEF. This is, in fact, the only mutation that shows no selectivity on its impact on the two RhoGEFs. This lack of selectivity and the 2-fold reduction in recruitment is likely due to the mutation disrupting the network of hydrogen bonds that is observed between R279 and both RhoGEFs. In the case of W233, it makes a small non-polar energy contribution exclusively to PDZRhoGEF. However, the W233A mutation involves a dramatic change in the physico-chemical properties of the residue which might introduce a conformational change that can affect interactions with p115RhoGEF more than interactions with PDZRhoGEF. When considering the T203A/D mutant, no receptor-promoted recruitment of p115RhoGEF could be observed. This lack of measurable recruitment did not result from a high constitutive recruitment since, in contrast with what is observed for the PDZRhoGEF, no constitutive recruitment could be detected for p115RhoGEF. In contrast, a sizable recruitment of PDZRhoGEF could be detected despite being blunted by the high constitutive recruitment. This is best illustrated by experiments in which increasing amounts of the TPα receptor were expressed (Fig. S8) revealing a robust receptor-promoted recruitment of PDZRhoGEF but not p115-RhoGEF at any receptor level tested. This reinforces the selective impact of the T203A/D mutations on p115RhoGEF. For two mutations (R200A, R200K) no receptor-promoted recruitment of either p115RhoGEF or PDZRhoGEF could be detected most likely as a result of the high constitutive recruitment described in Fig. 5A for these two mutants.

We identified mutations in both the Gα_13_ helical domain and in the GTPase domain that reduced Gα_13_ interactions with both p115RhoGEF and PDZRhoGEF (Fig. 5; Table S1). Mutations in most Sw I residues dramatically reduced recruitment of both RhoGEFs, with many mutations having a larger effect on interactions with p115RhoGEF. Exceptions were the Gα_13_ R201V and P202A substitutions that reduced interactions with p115RhoGEF by 26% and 37% (respectively) but had little effect on interactions with PDZRhoGEF, correlating with calculated contributions only to p115RhoGEF. Notably, T203A and T203D reduced p115RhoGEF and PDZRhoGEF recruitment by ∼84% and ∼95%, respectively.

As predicted by our computational analysis, nearly all Gα_13_ mutations reduced interactions with p115RhoGEF more than with PDZRhoGEF. Mutations in three helical domain residues reduced p115RhoGEF recruitment to the membrane by 27 to 81%, correlating with the predicted ΔΔG_total_ of these residues to p115RhoGEF, which was ∼4-7 kcal/mol. In contrast, PDZRhoGEF recruitment was significantly affected only by a double mutation in K94 and R97, which nevertheless also reduced PDZRhoGEF recruitment less than p115RhoGEF ( 71% vs 81%). Similar to mutations in the helical domain and Switch I, nearly all mutations in residues across Switch II, III, and the α3-β5 loop, which contribute more substantially to p115RhoGEF than to PDZRhoGEF, reduced Gα_13_ interactions with p115RhoGEF, while having little or no effect on interactions with PDZRhoGEF. Additionally, the few mutations that dramatically reduced recruitment to both RhoGEFs, such as in R230, E273, and R279, also showed substantial energy contributions to interactions with both RhoGEFs.

### Specific residues in the NTD and RH domains of p115RhoGEF are crucial for interactions with both Gα_13_ and Gα_12_

To quantify the contributions of p115RhoGEF residues to interactions Gα_13_ vs. Gα_12_, we mutated residues in the NTD and RH domain of p115RhoGEF (Fig. S9) and measured the effect on the recruitment to both G proteins using the GEMTA assay. To define the portion of the interaction that is due to NTD, we created a sensor with a scrambled NTD sequence (Fig. S9) which defines the residual signal resulting from an interaction due to the RH domain. Any point mutation in the NTD that would reduce the signal to the same level as the NTD-scrambled, would indicate a residue which is critical for the NTD-promoted interaction. NTD substitutions I23A/F and E27A reduced by ∼50% the TPαR-promoted p115RhoGEF binding to Gα_12_ and Gα_13_ bringing it to the level observed for the scrambled NTD sequence (Fig. 6A) indicating that these residues are critical for the NTD-promoted interaction.

**Figure 6.**
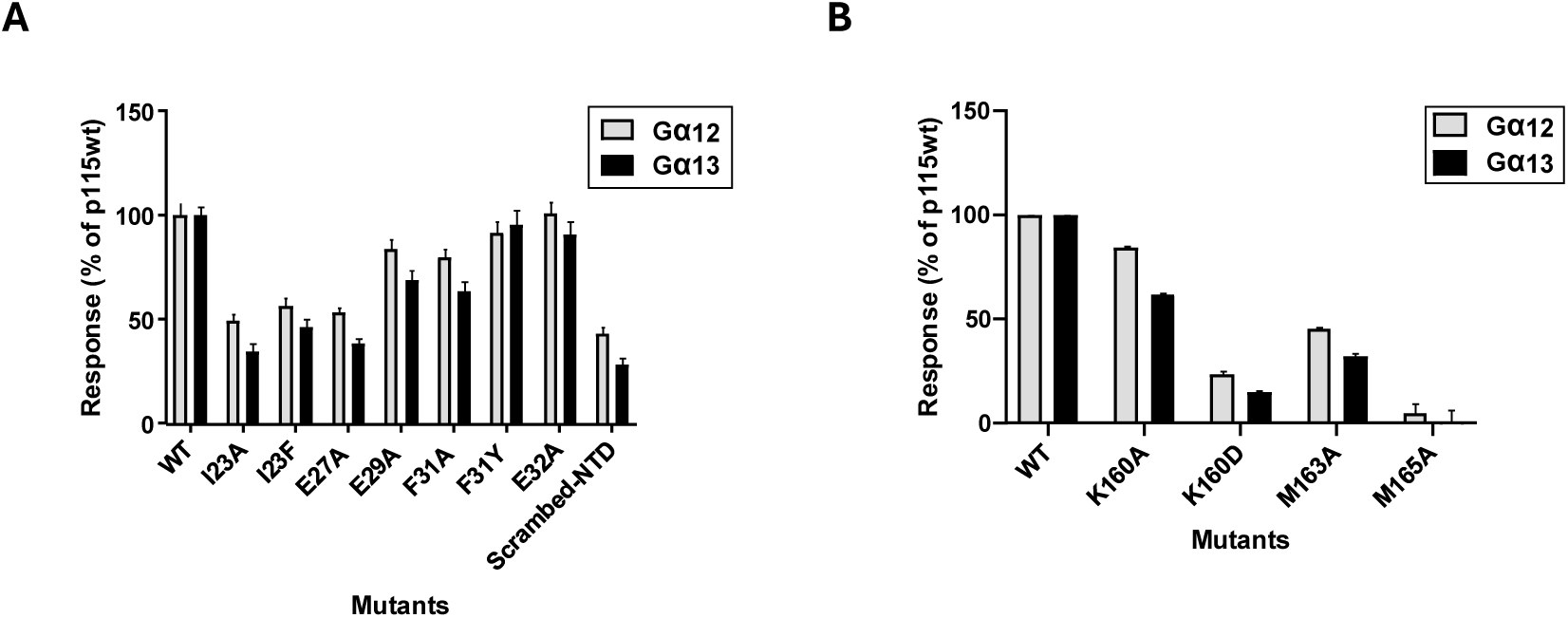
Effect of p115RhoGEF NTD and RH domain mutations on TPαR-promoted p115RhoGEF recruitment to Gα12 and Gα_13_. BRET was measured between WT or mutant forms of p115RhoGEF-Rluc biosensor (see supp. Fig S9) and the plasma-anchored rGFP-CAAX in the presence of heterologously expressed Gα12 (grey bars) or Gα_13_ (black bars) upon stimulation with the TPα agonist, U46619. The responses for mutations within the NTD and RH domains are illustrated in panels A and B, respectively, and expressed as % of the responses observed for the WT p115-RhoGEF sensor. In panel A, scrambled refers to a version of the biosensor in which the amino acid sequence of the NTD domain were scrambled; the remaining signal representing the contribution of the RH domain to the interaction.

In contrast, F31A/Y and E32A had little to no impact on p115RhoGEF binding to both Gα_12_ and Gα_13_ suggesting that their interactions with Gα_12/13_ is not essential for the recruitment. However, previous studies showed that F31 and E32 play a role in p115RhoGEF GAP activity, as mutations in these residues reduced such activity by at least 50% (31, 36), suggesting a specific role in mediating GAP activity but not binding affinity for the Gα subunits. RH mutations K160D, M163A, and M165A abolished the TPαR-induced recruitment of p115RhoGEF to both Gα_12_ and Gα_13_ (Fig. 6.B), correlating with our computational analysis as these residues make the highest calculated contributions from the RH domain to interactions with Gα_13_ (Fig. S6). Of notice, the K160A substitution had a milder effect than K160D, suggesting the charge reversal mutation plays a more significant role than merely disrupting interactions that rely on the positive charge of the lysine residue. Similar results were observed when using GPR35b, a GPCR that selectively activates Gα_13_ and not Gα_12_, upon stimulation with pamoic acid to promote the recruitment of p115RhoGEF to Gα_13_ (Fig. S10).

Taken together, these results indicate that the RH domain is essential for interactions, as demonstrated by the impact of the M165A mutation, whereas, although playing an important role, the NTD contributes to a lesser extent. There is a strong correlation between the effects of mutations in the N-terminal part of the NTD and the RH domain and our predictions of substantial contributions to interactions with Gα_13_.

## Discussion

In this study, we combined computational analysis with site-directed mutagenesis and ebBRET measurements to identify residues on both sides of the Gα_13_ interface with p115RhoGEF and PDZRhoGEF that affect these interactions. This combined analysis revealed that Gα_13_ contributions to p115RhoGEF are stronger than those to PDZRhoGEF, and that, accordingly, mutations in these Gα_13_ residues reduce p115RhoGEF recruitment more substantially than PDZRhoGEF recruitment. In a general view, our findings align with previous yet more limited biochemical studies, which tested only a few positions in Sw I, Sw II, and in the “effector binding site” – substantiating that the interactions of Gα_13_ with p115RhoGEF and PDZRhoGEF involve these regions. We demonstrated that the Gα helical domain also plays a substantial role in Gα_13_-p115RhoGEF interactions. Our combined approach suggests that the Gα_13_ Sw I region is a crucial element for recruitment of both p115RhoGEF and PDZRhoGEF. However, some residues from the Gα_13_ helical domain, Sw II, and Sw III were also essential for recruiting p115RhoGEF to activated Gα_13,_ as mutations in these positions completely abolished p115RhoGEF recruitment. We note that previous biochemical studies have shown that the Gα_13_ effector binding region, which we identified as needed for Gα_13_–p115/PDZRhoGEF interactions, is also central for p115RhoGEF GAP activity (33). Overall, our study provides a detailed understanding of how distinct Gα_13_ regions selectively influence the recruitment of p115RhoGEF and PDZRhoGEF and highlighting individual residues that differentially modulate interactions with these RhoGEFs, thereby contributing to the specificity and regulation of Gα_13_ signaling pathways.

Looking at the p115RhoGEF side of the interface, our results demonstrated that the NTD is needed not only for GAP activity (36), but also that it is necessary for high-affinity recruitment of Gα_12/12_. In addition, our findings that mutations across the RH domain reduced recruitment to Gα_12/13_ around 2-fold, revealed its role mediating interactions with Gα_12/13_, adding to its previously reported role in GAP activity (25). Looking forward, our detailed residue-level mapping of specific interaction determinants among these proteins lays the groundwork for the rational design of new therapeutic agents targeting Gα_13_ interactions with p115RhoGEF and PDZRhoGEF.

## Acknowledgements

This work was supported by the Israel Science Foundation (grant 1454/13) to MK, grants from the Canadian Institutes of Health Research (CIHR: PJT-183758) and the Natural Science and Engineering Research Council of Canada (NSRC: RGPIN-2019-05556) to MB, the International Development Research Centre (IDRC), the Israel Science Foundation (ISF), and the Azrieli Foundation (grant 3512/19) to MK and MB, and by a grant from the Council for Higher Education through the Data Science Research Center at the University of Haifa to MK.

## Author contributions

AB designed and performed experiments, analyzed data and wrote the manuscript, VL designed and performed experiments and revised the manuscript, CL designed and performed experiments, supervised research and revised the manuscript, MB designed experiments, supervised research and wrote the manuscript, MK designed experiments, analyzed data, supervised research, and wrote the manuscript.

## MATERIALS AND METHODS

### Protein structures

The following representative 3D structures were used in our analysis and visualization of Gα_13_ subunits with different partners (with PDB IDs): Gα_13/i1_–p115RhoGEF bound to GDP/AlF_4_ (1SHZ) (31), Gα_13_–p115RhoGEF bound to GDP/AlF_4_ (3AB3) (33), Gα_13_–PDZRhoGEF bound to GDP/AlF_4_ (3CX7), and Gα_13_–PDZRhoGEF bound to GTPγS (3CX8) (32). Missing short segments in the following PDB entries were modeled using Loopy (49) and partial or missing side chains were modeled using Scap (49): 1SHZ (p115RhoGEF residues 38-43, 87-92, and 123-132), 3AB3 (Gα_13_ residues 337-340 and p115RhoGEF residues 35-45, 86-92, and 122-133), 3CX7 (Gα_13_ residues 337-340 and PDZRhoGEF residues 319-321), 3CX8 (Gα_13_ residues 337-340 and PDZRhoGEF residues 319-321). Hydrogen atoms were added using CHARMM, and the structures were subjected to conjugate gradient minimization with a harmonic restraint force of 50 kcal/mol/Å^2^ applied to the heavy atoms. 3D structural visualization was carried out with the PyMOL molecular graphics program (https://www.pymol.org/).

### Energy calculations to map residue-level specificity determinants

We followed the methodology described previously (50-53) to analyze the per-residue contributions of Gα_13_ residues to interactions with p115RhoGEF and PDZRhoGEF in the crystal structures mentioned above. The Finite Difference Poisson–Boltzmann (FDPB) method, as implemented in DelPhi (54), was used to calculate the net electrostatic and polar contributions (ΔΔG_elec_) of each residue within 15Å of the dimer interface. For each residue, electrostatic contributions from the side chain and/or originating from the main chain were calculated separately. Residues contributing ΔΔG_elec_ ≥ 1 kcal/mol to the interactions (twice the maximal numerical error of the electrostatic calculations, see (44)) were deemed as substantially contributing to the interaction (50). Non-polar energy contributions (ΔΔG_np_) were calculated as a surface-area proportional term by multiplying the per-residue surface area buried upon complex formation, calculated using Surfv (55), by a surface tension constant of 0.05 kcal/mol/Å^2^ (44, 50). Residues contributing ΔΔG_np_ ≥ 0.5 kcal/mol to the interactions (namely, those that bury more than 10Å^2^ of each protein surface upon complex formation) were defined as making substantial non-polar contributions (52, 53). To reduce false positives and negatives, we applied a consensus approach across comparable biological replicates in multiple PDB structures (1SHZ, 3AB3, 3CX7, 3CX8), substantially improving prediction accuracy or across multiple dimers in an asymmetric unit (see Fig. S1 and S2). Analyzing the two Gα_13_-PDZRhoGEF complexes, with GTPγS (3CX8) and GDP/AlF_4_ (3CX7), we combined the results from both structures into a consensus approach, after visually validating that each individual contributions observed only in one structure had a plausible origin across the complex.

### Reagents

Dulbecco’s phosphate-buffered saline (D-PBS), Dulbecco’s modified Eagle’s medium (DMEM), Trypsin, PBS, penicillin/streptomycin, fetal bovine serum, and newborn calf serum were from Wisent Bioproducts. Polyethylenimine (PEI) was purchased from Alfa Aesar (Thermo Fisher Scientific). U46619, and pamoic acid were from Cayman Chemical. Prolume Purple and Coelenterazine 400a were purchased from Nanolight Technologies. Restriction enzymes and Q5 mutagenesis kit were purchased at New England Biolabs.

### Plasmid DNA Constructs

The following protein constructs were described previously: HA-TPαR (56), GRK2-D110A-GFP10 and RlucII-Gγ5 (47), rGFP-CAAX (43), and PDZ RhoGEF-RlucII sensor (57). Gα_12_, Gα_13_ and Gβ1 were purchased from cDNA.org (Bloomsburg University). The construct encoding a codon-optimized GPR35b was synthetized in pCDNA3.1 Zeo(+) at GeneArt (ThermoFisher). The p115 RhoGEF (1-244)-RlucII was created from a DNA fragment synthetized and subcloning into NheI + HindIII sites of pCDNA3.1 Hygro (+) GFP10-RlucII, replacing the GFP10 coding sequence. The p115 RhoGEF (21-244)-RlucII construct, used in this study, was created by a deletion of the first 20 codons of p115 RhoGEF (1-244)-RlucII, using the Q5 mutagenesis kit. Substitutions in Gα_13_ and p115 RhoGEF (21-244)-RlucII were created using the Q5 mutagenesis kit with oligonucleotides synthetized at ThermoFisher.

### Cell Culture and Transfection

HEK 293SL cells (43) were propagated in plastic flasks and grown at 37 °C in 5% CO_2_ and 90% humidity. Cells (3.2 × 10^4^ cells in 1 mL) were transfected in suspension with 1 μg of DNA (plasmid DNA and adjusted with ssDNA), complexed with linear PEI (molecular weight 25,000, 3:1 PEI:DNA ratio) and seeded (3.2 × 10^4^ cells/well) in poly-D-Lysine coated white 96-well plates.

### p115 RhoGEF and PDZRhoGEF translocation assay

For receptor-mediated Gα_13_ & Gα_12_ activation measured by p115 RhoGEF and PDZ RhoGEF recruitment, HEK 293SL cells were transfected with pCDNA3.2 plasmids encoding receptors HA-TPαR: or GPR35b: along with Gα_12_, or Gα_13_, rGFP-CAAX with either p115 RhoGEF-RlucII or PDZ RhoGEF-RlucII. After a 48-h incubation, cells were washed once with PBS, then incubated in Tyrode’s buffer, at 37C for one hour. Cells were then exposed to either receptor agonists (U46619 for TPαR or pamoic acid for GPR35b) or vehicle for 10 min. Prolume Purple (RlucII substrate; 1 μM final) is added at 4min of stimulation, for a total incubation of 10min prior to reading BRET.

### Gβ1/RlucII-Gγ5 release assay

For receptor-mediated G13 activation measured by Gβγ release from Gα as captured by GRK2-D110A-GFP10, HEK 293SL cells were transfected with pCDNa3 plasmids expressing HA-TPαR, Gα_13_, Gβ1, RlucII-Gγ5 and GRK2-D110A-GFP10. After a 48-h incubation, cells were washed once with PBS, then incubated in Tyrode’s buffer, at 37C for an hour. Cells were then exposed to either the TP agonist U46619 or vehicle (Methyl Acetate at 0.1% final) for 10 min. Coelenterazine 400a (RlucII substrate; 2.5 μM final) is added at 5min of stimulation, for a total incubation of 10min prior to reading BRET.

### BRET measurements and analysis

Plates were read on a Tristar2 LB 942 from Berthold Technologies equipped with filters for BRET2 (donor: 400/70 nm and acceptor: 515/20 nm), for detecting the RlucII (donor) and rGFP (acceptor) light emissions, respectively. Data and statistical analysis were performed using GraphPad Prism 10.4.1 (GraphPad Software). Quantitative data are expressed as the mean, and error bars represent the SEM, unless otherwise indicated. Curve fitting was performed using four parameters nonlinear regression.

