## Supplementary Figures 1-10, Supplementary Table 1 for "Molecular determinants underlying differential recruitment of p115RhoGEF and PDZRhoGEF to activated Gα_13_"

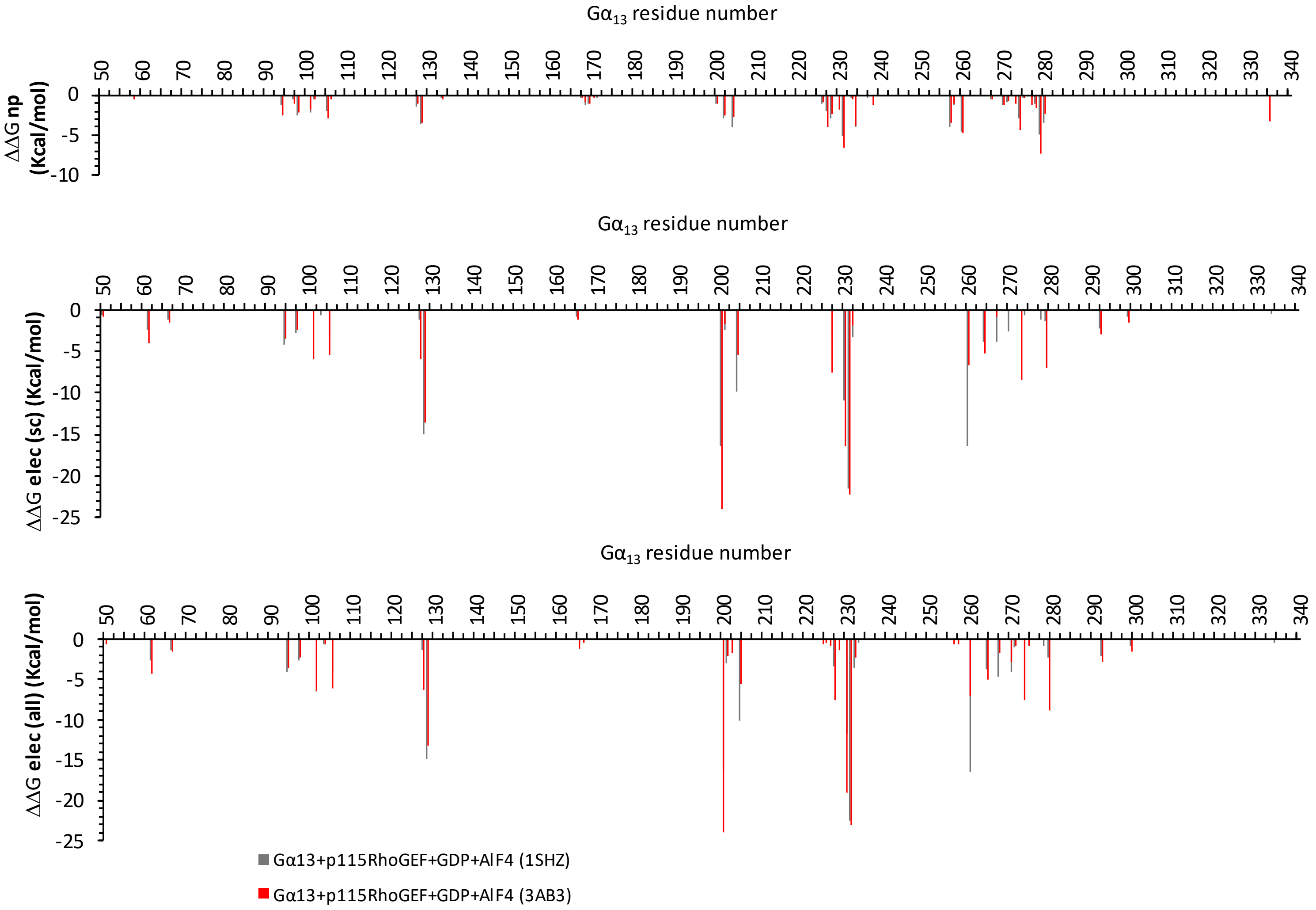


**Figure S1. Comparing Gα$_{13}$ per-residue energy contribution to interactions with p115RhoGEF.** Panels show the results across two comparable structures: Gα$_{13}$/Gα$_i$-p115RhoGEF with GDP/AlF$_4$ (PDB ID 1SHZ) and Gα$_{13}$-p115RhoGEF with GDP/AlF$_4$ (PDB ID 3AB3). Top panel shows non-polar (np) energy contributions, middle panel shows electrostatic contributions from the side-chain (ΔΔG$_{elec}$ (sc)), and the bottom panel shows electrostatic contributions from the entire residue (ΔΔG$_{elec}$ (all)), calculated as described in Methods. When comparing the two complexes we considered the Gα$_{13}$-p115RhoGEF complex (3AB3) with the wild-type protein and the better resolution as the more representative of the two structures.

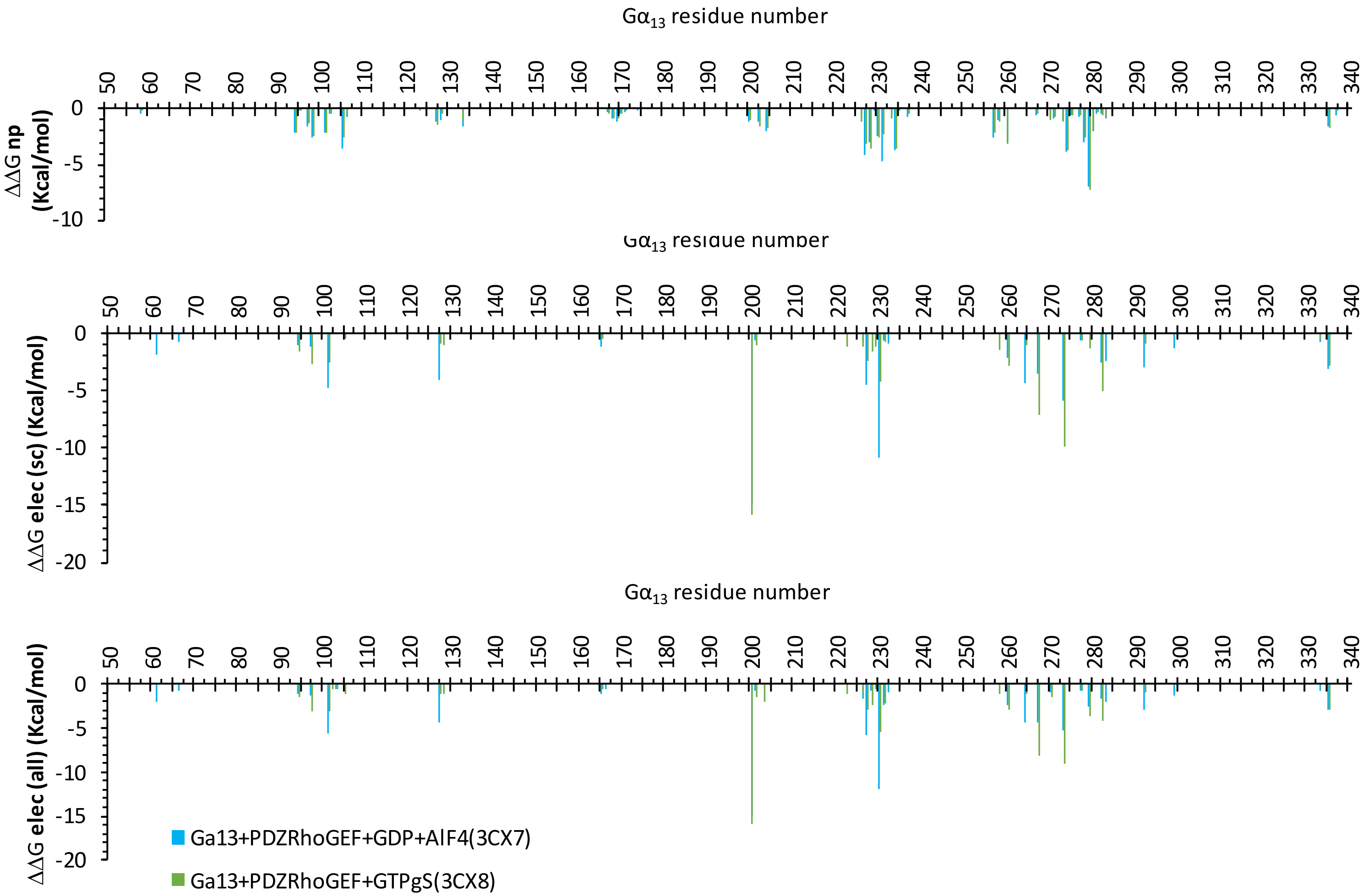


**Figure S2. Comparing Gα$_{13}$ per-residue energy contribution to interactions with PDZRhoGEF.** Panels show the results across two comparable structures: Gα$_{13}$-PDZRhoGEF with GDP/$AlF_4$ (PDB ID 3CX7) and Gα$_{13}$-PDZRhoGEF with GTPγS (PDB ID 3CX8). Top panel shows non-polar (np) energy contributions, middle panel shows electrostatic contributions from the side-chain ($\Delta\Delta G_{elec}$ (sc)), and the bottom panel shows electrostatic contributions from the entire residue ($\Delta\Delta G_{elec}$ (all)), calculated as described in Methods. When comparing the two complexes we considered the Gα$_{13}$-PDZRhoGEF with GDP/$AlF_4$ complex (3CX7) as the more representative structure.

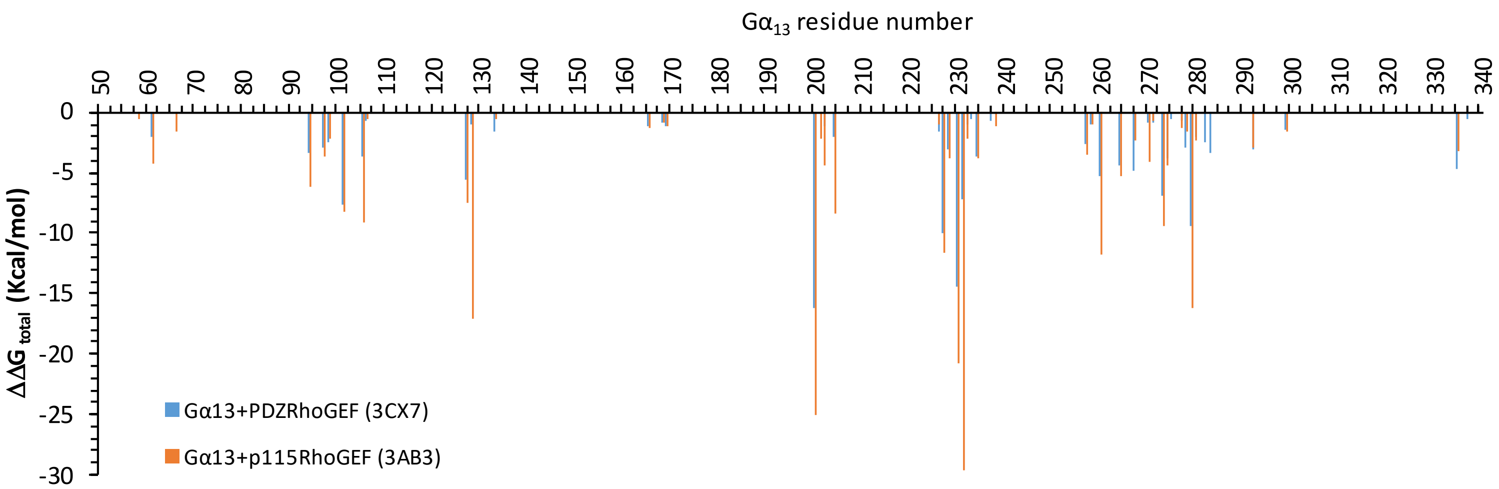


**Figure S3. Total G$\alpha_{13}$ per-residue energy contributions to interactions with p115RhoGEF and PDZRhoGEF.** $\Delta\Delta G_{total}$ equals to the sum of $\Delta\Delta G_{np}$ and $\Delta\Delta G_{elec}$ (all) for the interactions of G$\alpha_{13}$ with each partner, as in Figs. S1 and S2.

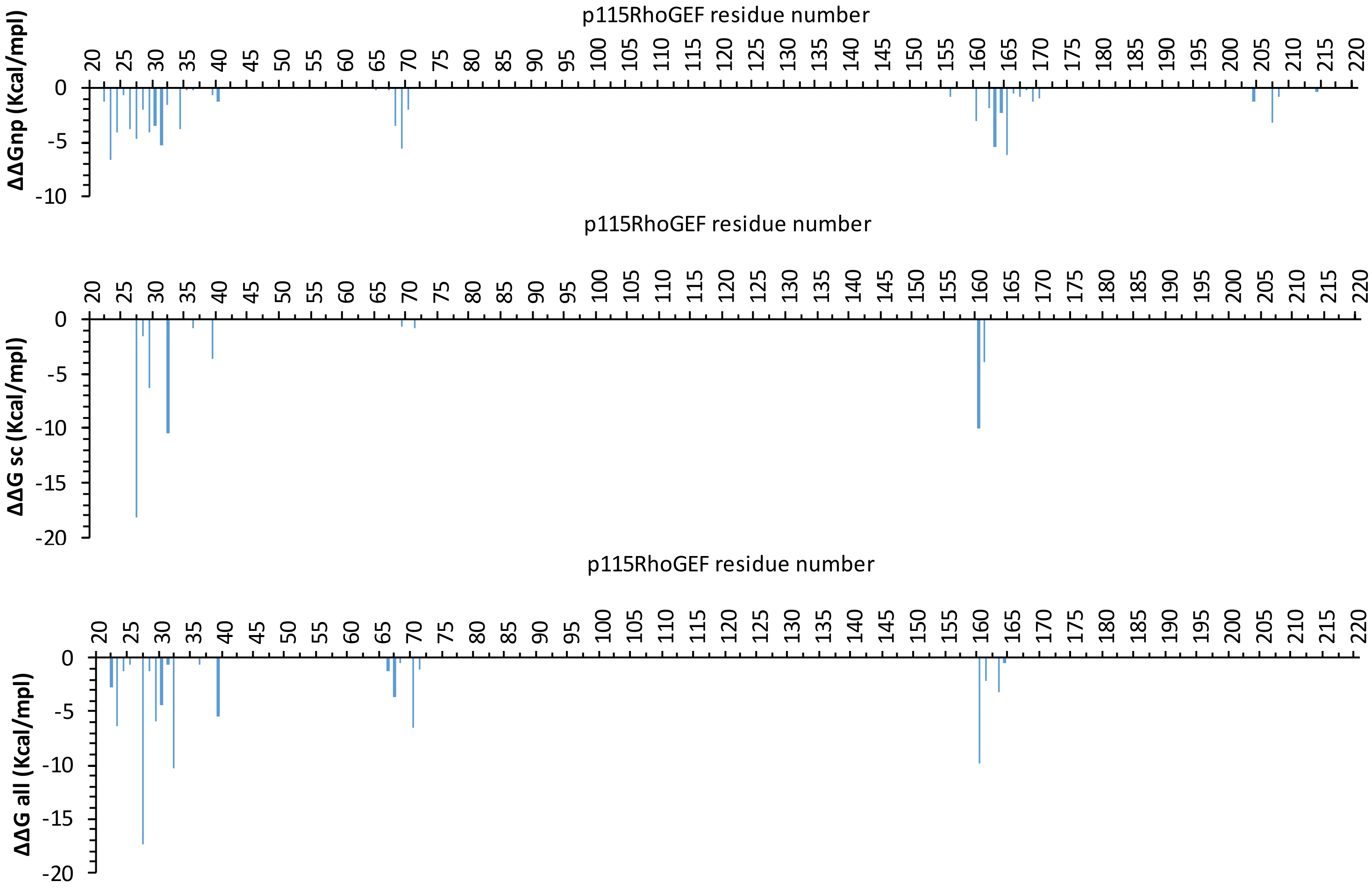


**Figure S4. p115RhoGEF per-residue energy contributions to interactions with $G\alpha_{13}$.** Panels show the energy results for non-polar (np) contributions, electrostatic contributions from the side-chain ($\Delta\Delta G_{elec}$ (sc)), and electrostatic contributions from the entire residue ($\Delta\Delta G_{elec}$ (all)), as above.

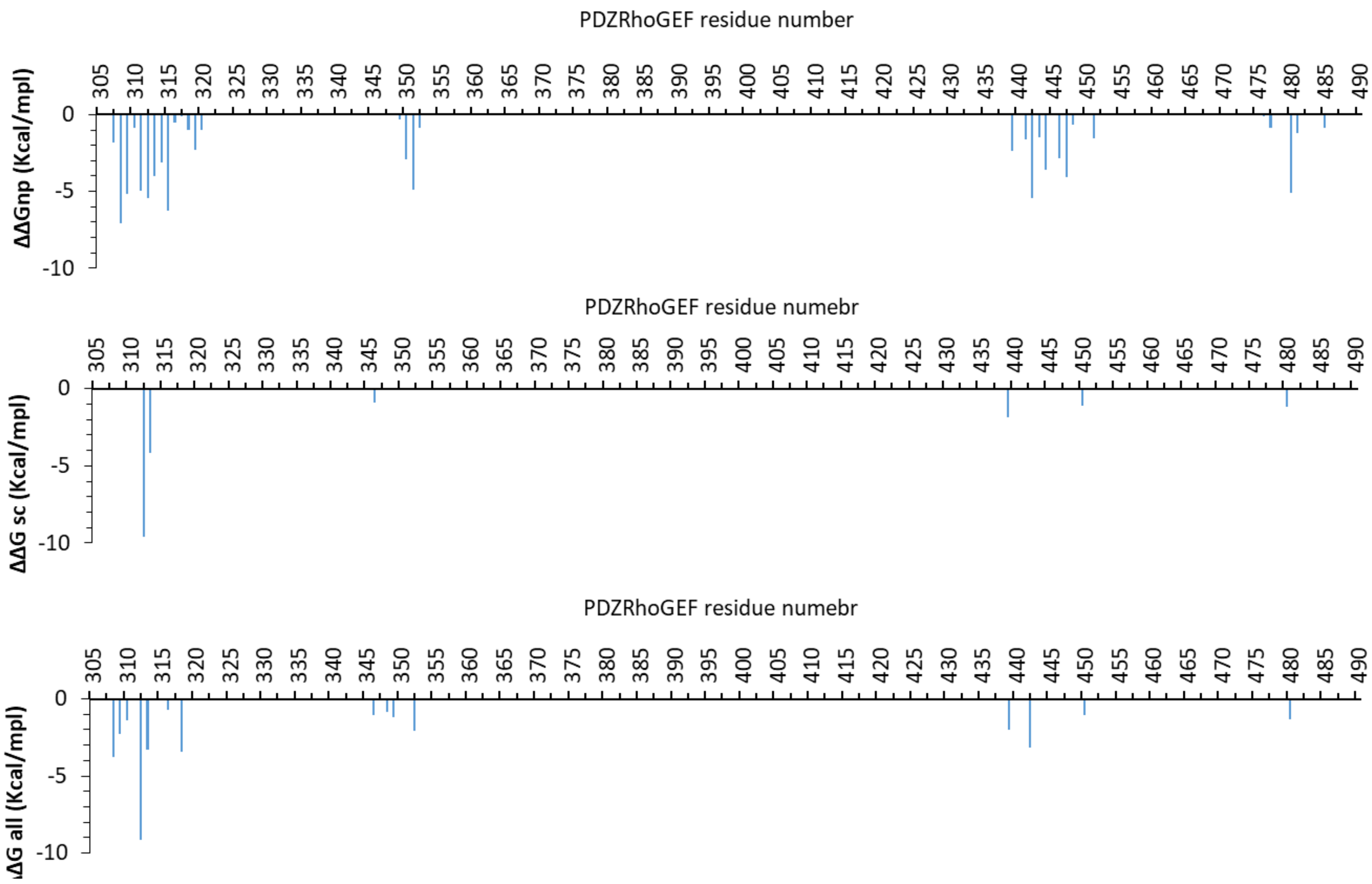


**Figure S5. PDZRhoGEF per-residue energy contributions to interactions with G$\alpha_{13}$.** Panels show the energy results for non-polar (np) contributions, electrostatic contributions from the side-chain ($\Delta\Delta G_{elec}$ (sc)), and electrostatic contributions from the entire residue ($\Delta\Delta G_{elec}$ (all)), calculated as above.

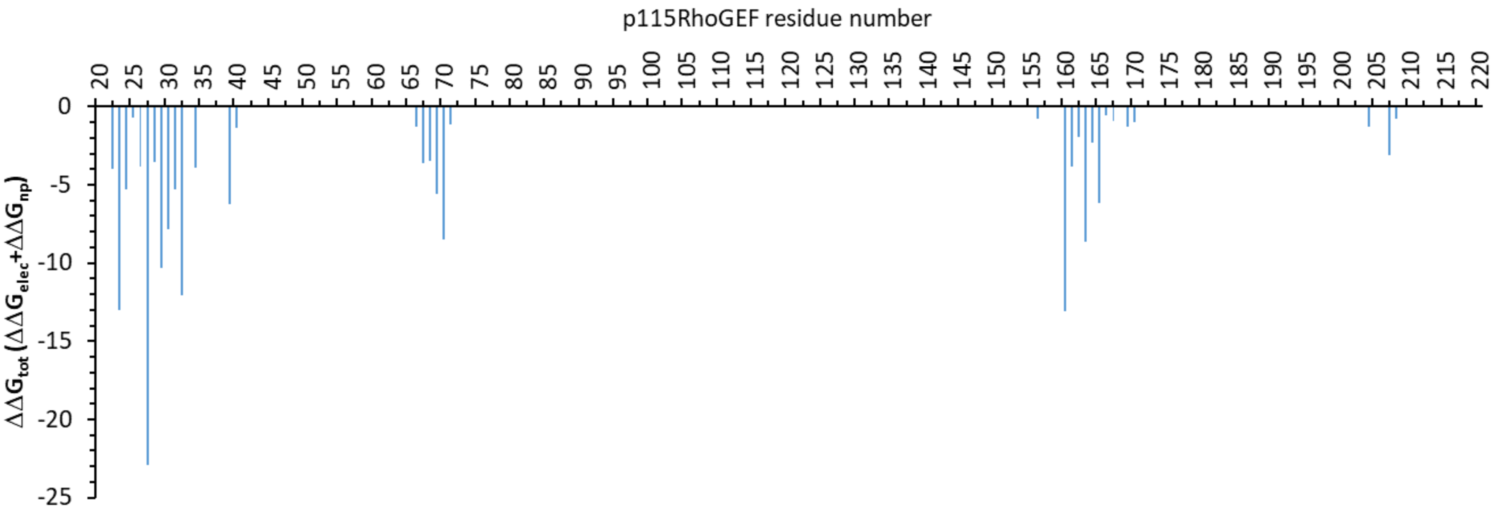


**Figure S6. Total p115RhoGEF per-residue energy contributions to interactions with G$\alpha_{13}$.** $\Delta\Delta G_{total}$ equals to the sum of $\Delta\Delta G_{np}$ and $\Delta\Delta G_{elec}$ (all) for the interactions of G$\alpha_{13}$ with each partner, as in Fig. S3.

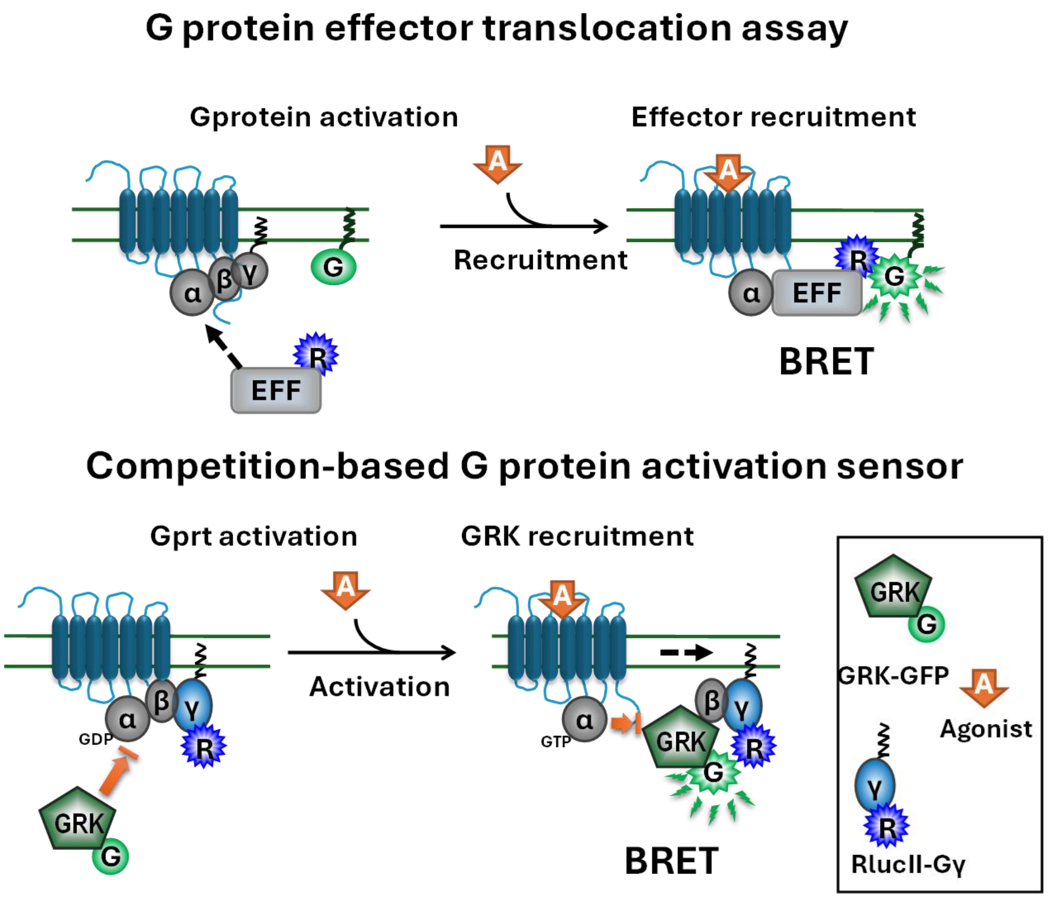


**Figure S7**. **Illustration of the BRET-based sensor systems used. Top panel:** G protein effector membrane translocation assay (GEMTA). BRET is measured between a G protein subtype-specific effector (ex: p115RhoGEF or PDZRhoGEF for $G\alpha_{12/13}$) fused to Rluc and rGFP anchored at the plasma membrane via a CAAX-box sequence. The recruitment of the Rluc-effector to the activated $G\alpha$ subunit bring the Rluc in close proximity of the GFP resulting in an increase in BRET. **Bottom panel:** GRK-$G\beta\gamma$-based assay. BRET is measured between GRK-GFP and $\beta\gamma$-Rluc in the presence of heterologously expressed $G\alpha$. Sensors and agonist are depicted as in the caption. Upon activation of the $G\alpha\beta\gamma$ heterotrimeric G protein, GRK-GFP is recruited to the released $G\beta\gamma$-Rluc resulting in an increase in BRET.

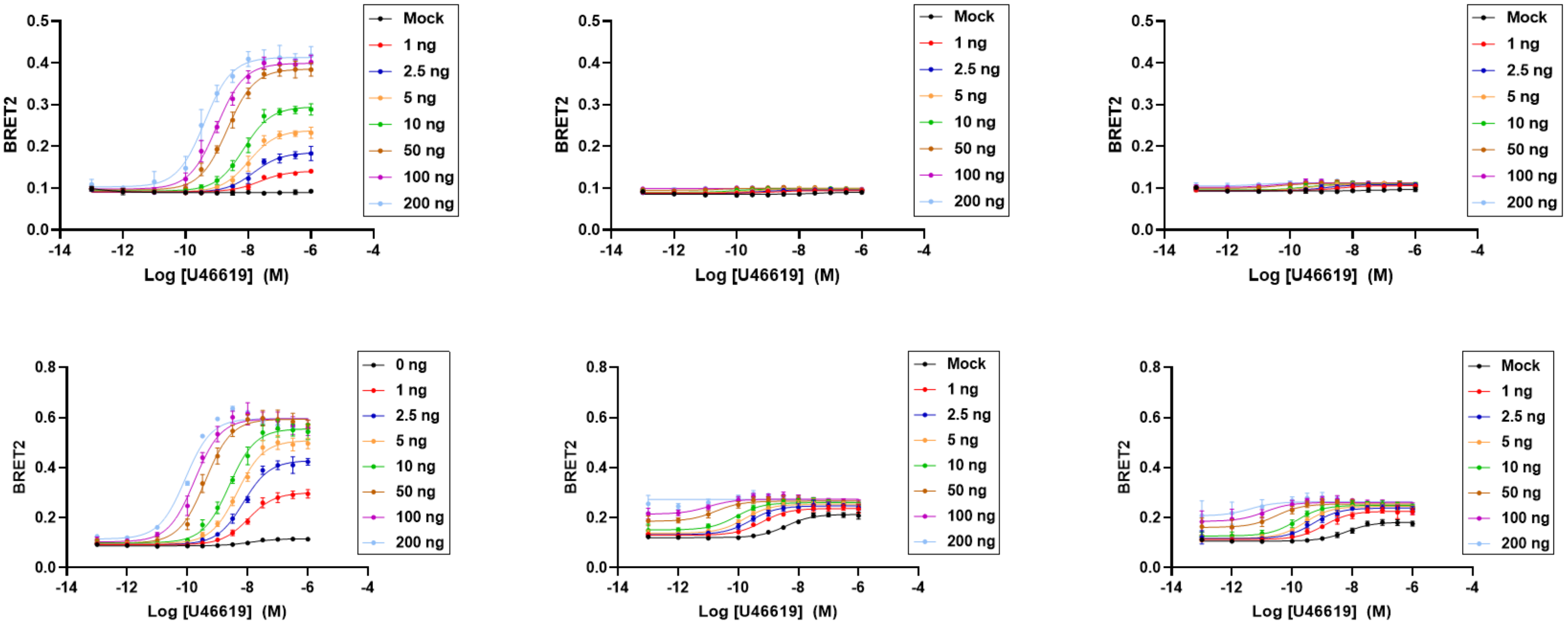


**Figure S8. Effect of T203A/D substitutions in $G\alpha_{13}$ on the TPα-promoted recruitment of p115RhoGEF to $G\alpha_{13}$ assessed using GEMTA sensor.** BRET was measured between the plasma-anchored rGFP-CAAX and p115RhoGEF-Rluc (top row) or PDZRhoGEF-Rluc (bottom row) in the presence of heterologously expressed WT (top and bottom left panels) or T203A (top and bottom middle panels) or T293D (top and bottom right panels) mutant forms of $G\alpha_{13}$ or following activation with increasing concentration of the thromboxane receptor TPα agonist U46619. The experiments were carried out in the presence of increasing concentration of the TPαR expressing plasmid (0-200 ng).

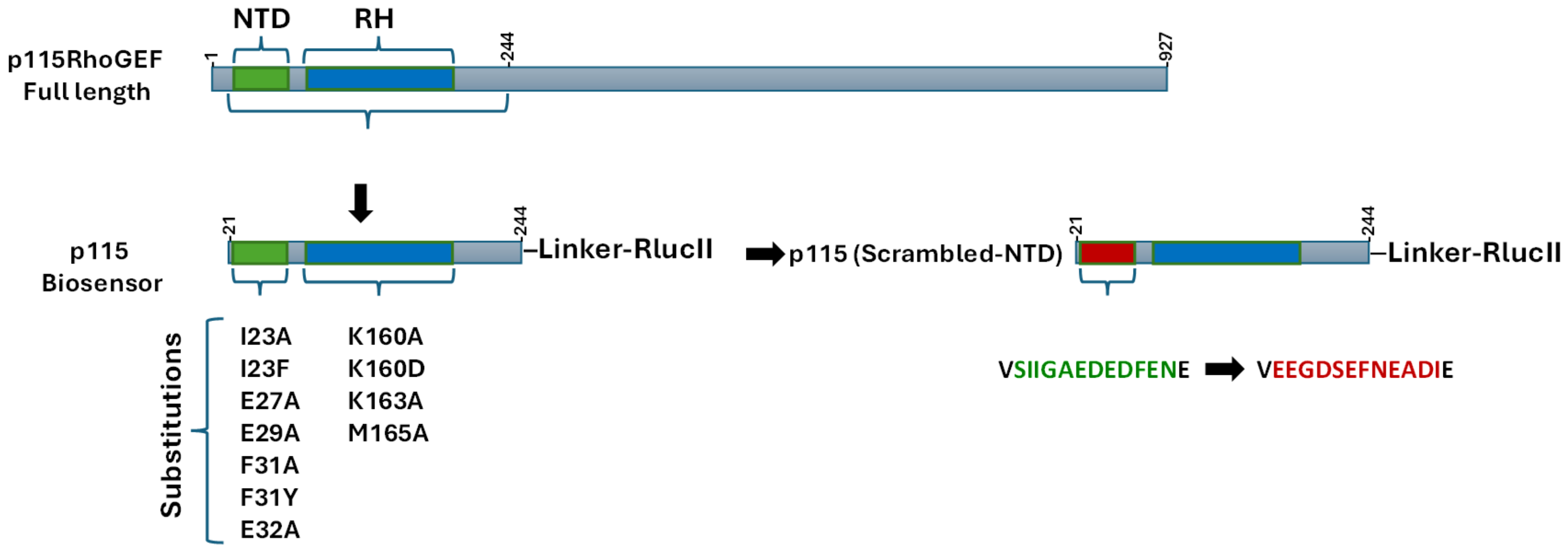


**Figure S9. Illustration of the p115RhoGEF biosensor. Top panel:** Schematic representation of the full length p115RhoGEF sequence highlighting the NTD and RH domains. **Bottom panel:** Schematic representation of the p115RhoGEF fragment (21-244) that was fused to Rluc to generate the p115RhoGEF-Rluc biosensor. This fragment incudes the two domains (NTD and RH) responsible for the interaction with $G\alpha_{12/13}$. The illustration also indicates the positions of the NTD and RH domain mutations studied. The WT (green) and scrambled (red) sequences of the NTD that were used to assess the global contribution of the RH domain are also shown.

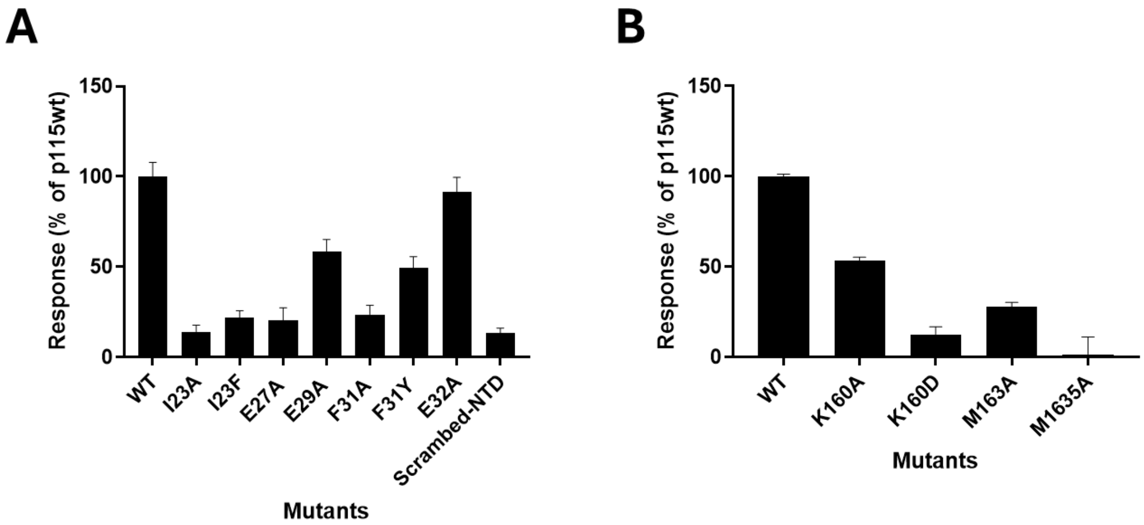


**Figure S10. Effect of p115RhoGEF NTD and RH domain mutations on GPR35b-promoted p115RhoGEF recruitment to G**$\boldsymbol{\alpha}_{13}$**:** BRET was measured between WT or mutant forms of p115-RhoGEF-Rluc biosensor (see supp. Fig. S9) and the plasma-anchored rGFP-CAAX in the presence of heterologously expressed G$\alpha_{13}$ upon stimulation with the GPR35b agonist, pamoic acid. The responses for mutations within the NTD and RH domains are illustrated in panels A and B, respectively, and expressed as % of the responses observed for the WT p115RhoGEF sensor. In panel A, scrambled refers to a version of the biosensor in which the amino acid sequence of the NTD domain were scrambled to establish the contribution of the RH domain to the interaction.

| TPaR/U46619 | p115 | PDZ | p115 | PDZ |
|---|---|---|---|---|
| G13 mutants | Span in % | Span in % | pEC50 | pEC50 |
| WT | 99 | 98 | 8.4 | 8.9 |
| K94A | 51 | 79 | 8.7 | 8.8 |
| R97E | 28 | 67 | 9.2 | 9.3 |
| R97Q | 42 | 92 | 8.8 | 9.1 |
| K94E/R97E | 19 | 29 | 9.7 | 9.6 |
| K105A | 62 | 86 | 8.6 | 8.7 |
| T127E | 44 | 98 | 8.8 | 9.2 |
| R128A | 63 | 106 | 8.8 | 9.0 |
| F168A | 49 | 67 | 8.5 | 8.6 |
| R200A | 3 | 8 | 10.6 | 9.9 |
| R200K | -3 | 8 | 11.0 | 10.5 |
| R201V | 74 | 106 | 8.9 | 9.7 |
| P202A | 63 | 97 | 8.9 | 9.8 |
| T203A | 3 | 23 | 10.0 | 10.1 |
| T203D | 7 | 26 | 9.6 | 9.9 |
| K204A | 28 | 91 | 9.4 | 9.5 |
| R230A | 10 | 30 | 8.8 | 8.5 |
| K231A | 65 | 89 | 8.2 | 8.5 |
| K231Q | 72 | 102 | 8.4 | 8.6 |
| W233A | 3 | 20 | 8.7 | 9.3 |
| M257A | 101 | 108 | 8.7 | 8.8 |
| R260A | 47 | 98 | 8.6 | 8.8 |
| E273R | 8 | 60 | 8.8 | 9.3 |
| R279A | 57 | 68 | 8.6 | 9.0 |
| R279D | 4 | 29 | 9.4 | 9.5 |
| V280W | 115 | 98 | 8.4 | 9.1 |
| S282A/N283A | 70 | 95 | 8.4 | 8.5 |
| Q337A | 94 | 103 | 8.4 | 8.7 |

**Table S1. Effect of G$\alpha_{13}$ mutations on TP$\alpha$R-promoted p115RhoGEF and PDZRhoGEF recruitment.** Concentration response curves for the TP$\alpha$R-promoted p115RhoGEP-Rluc and PDZRhoGEF-Rluc/rGFP-CAAX BRET response were performed for the WT and the indicated mutant forms of G$\alpha_{13}$. The maximal BRET response (span) observed are expressed as % of the asymptotic maximal response obtained to the WT G$\alpha_{13}$. The potency of the responses is expressed as pEC50. The heat-map of the values goes from red (the lowest) to green (the highest).
